# A periplasmic regulator establishes adaptive impermeability to control carbapenem entry

**DOI:** 10.64898/2026.08.14.744805

**Authors:** Verena Ducret, Catarina Gonçalves Milho, Karl Perron

## Abstract

Outer-membrane permeability is a major determinant of antibiotic susceptibility in Gram-negative bacteria and is generally thought to be controlled through transcriptional regulation of porin expression. Here we identify the small periplasmic protein PtrA as a regulator of OprD-dependent carbapenem permeability in *Pseudomonas aeruginosa*. PtrA promotes imipenem resistance without altering OprD abundance, associates with OprD-containing membrane complexes and reduces intracellular imipenem entry. Using zinc and copper as complementary physiological signals, we show that PtrA-mediated permeability control is mechanistically distinct from CzcRS-dependent repression of *oprD* and precedes transcriptional porin depletion. These findings define a two-phase mechanism in which rapid periplasmic regulation provides an immediate adaptive response before transcriptional remodeling of the outer membrane. Our work identifies adaptive impermeability as a previously unrecognized mechanism linking environmental sensing to dynamic control of bacterial outer-membrane permeability.

## INTRODUCTION

Carbapenem resistance in *Pseudomonas aeruginosa* remains a major clinical challenge and is frequently driven by reduced outer-membrane permeability, most commonly through loss or repression of the porin OprD, the principal uptake pathway for imipenem (Do Rego and Timsit 2023). While the transcriptional regulation of *oprD* has been extensively characterized, considerably less is known about how porin activity itself can be rapidly modulated in response to acute environmental stress.

Outer-membrane permeability is highly dynamic and continuously adjusted to balance nutrient acquisition with protection against toxic compounds. Known adaptive mechanisms rely on transcriptional remodeling of porin expression, changes in porin composition, or physicochemical modulation of pore conductance in response to environmental cues such as osmolarity, pH, metabolic state or ionic conditions (Nikaido 2003; Delcour 2009). However, these mechanisms primarily alter porin abundance or channel properties rather than rapidly regulating the activity of already assembled porins. Whether bacteria possess dedicated periplasmic factors capable of transiently reducing porin permeability before transcriptional adaptation occurs remains unknown. Rapid regulation of porin permeability would be particularly advantageous during sudden environmental transitions, such as those encountered upon host entry or phagocytosis, where immediate protection may be required before transcriptional adaptation can occur.

Consistent with this hypothesis, the bacterial periplasm is ideally positioned to integrate environmental signals. During infection, phagocytic cells expose bacteria to elevated concentrations of Zn(II) and Cu(I)/Cu(II) within the phagolysosome (Sheldon and Skaar 2019). In *P. aeruginosa*, excess Zn and Cu activate the CzcRS and CopRS two-component systems, respectively, triggering adaptive responses that coordinate metal detoxification, envelope remodeling, and antibiotic resistance (Ducret et al. 2020). In particular, activation of the Zn-responsive CzcRS pathway induces metal efflux systems while repressing *oprD* expression, thereby linking Zn stress to imipenem resistance (Perron et al. 2004; Caille et al. 2007).

However, although *oprD* transcription is rapidly repressed following Zn or Cu exposure, OprD itself does not immediately disappear from the outer membrane. Like many β-barrel porins, OprD is highly stable once assembled, exhibiting slow turnover and limited proteolytic degradation (Haltia and Freire 1995; Hartojo and Doyle 2024). Consequently, OprD molecules remain detectable for several hours following transcriptional shutdown (Ducret et al. 2021). Together, these observations led us to hypothesize that metal-induced periplasmic proteins transiently regulate OprD permeability before transcriptional remodeling of the outer membrane becomes effective. Among the proteins induced during Zn and Cu stress, the small periplasmic protein PtrA (PA2808) emerged as a compelling candidate. Although previously implicated in copper tolerance (Elsen et al. 2011), its physiological function remains unknown.

Here we identify PtrA as the first component of a previously unrecognized mechanism of adaptive impermeability in *P. aeruginosa*. PtrA transiently reduces OprD-dependent carbapenem permeability without altering OprD abundance and acts before the well-established CzcRS-dependent transcriptional repression of *oprD*. Together, these findings uncover a two-phase mechanism coupling metal sensing to rapid regulation of outer-membrane permeability before transcriptional remodeling becomes effective.

## RESULTS

### PtrA promotes imipenem resistance without altering OprD abundance

Because Zn/Cu-induced carbapenem resistance has long been attributed to transcriptional repression of *oprD* (Perron et al. 2004), we asked whether the Zn/Cu-responsive periplasmic protein PtrA contributes to this adaptive response. IPTG-induced expression of *ptrA* was sufficient to confer a marked increase in imipenem resistance even in the absence of metal (Fig. 1A). Importantly, the C-terminally 3×FLAG-tagged PtrA construct (PtrA-3F) fully recapitulated the resistance phenotype of the untagged protein, demonstrating that the C-terminal tag does not impair PtrA function and validating its use throughout this study. Unexpectedly, OprD remained readily detectable after both 5 h and 24 h of PtrA induction (Fig. 1B), demonstrating that PtrA-mediated resistance occurs without detectable loss of the porin. We next asked whether PtrA indirectly reprograms genes involved in β-lactam resistance (Torrens et al. 2019; Zahedi bialvaei et al. 2021; Amisano et al. 2025). qRT-PCR analysis revealed no significant changes in the expression of *oprD*, *opdP*, *oprM*, *oprN* or *ampC* following PtrA induction (Fig. 1C). Thus, neither altered porin expression, activation of efflux systems, nor increased β-lactamase expression accounted for the observed phenotype. Together, these findings demonstrate that PtrA promotes imipenem resistance through a mechanism distinct from the established transcriptional pathways controlling carbapenem susceptibility.

**Figure 1.**
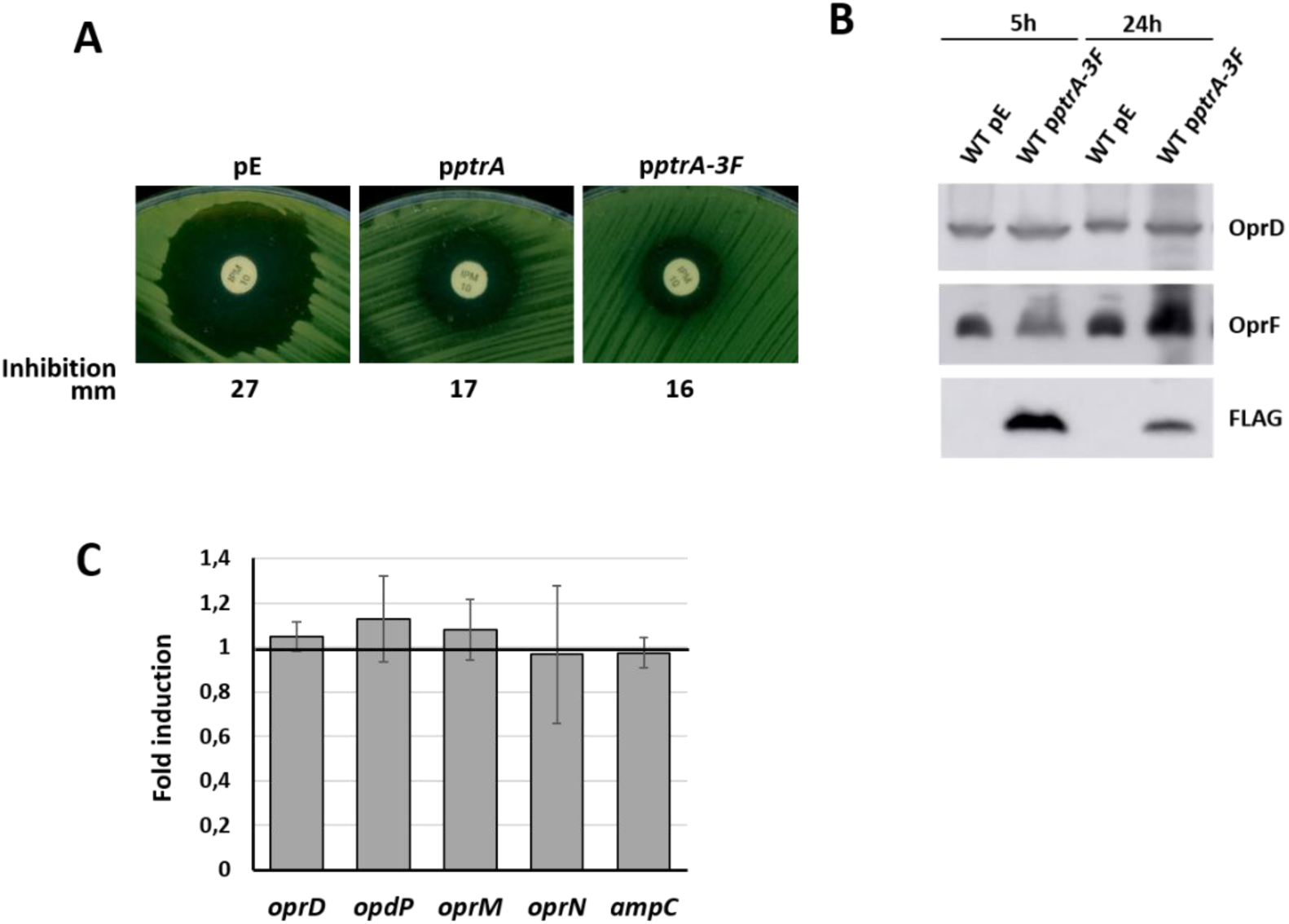
PtrA overexpression confers imipenem resistance. **(A)** Representative imipenem antibiograms of the WT strain carrying the empty vector (pE), *ptrA* (p*ptrA*), or the C-terminally 3×FLAG-tagged *ptrA* construct (p*ptrA*-*3F*). Assays were performed on LB agar supplemented with 1 mM IPTG. Inhibition zone diameters (mm) are indicated below each image. **(B)** Western blot analysis of OprD following 5 h or 24 h of PtrA overexpression (1 mM IPTG). **(C)** qRT-PCR analysis of selected genes potentially involved in β-lactam resistance following *ptrA* overexpression in the WT strain (WT p*ptrA*) induced with 1 mM IPTG. Relative transcript levels were compared to those of the WT strain carrying the empty vector (WT pMMB66EH), whose expression level was arbitrarily set to 1. Data represent the mean ± SD of three independent biological replicates. Statistical significance was assessed using two-tailed unpaired Student’s *t*-tests; all comparisons were not significant (*P* > 0.05).

To determine whether PtrA broadly affects envelope permeability or specifically alters carbapenem susceptibility, we compared the activity of several antibiotic classes (Table 1). PtrA overexpression selectively reduced susceptibility to imipenem and, to a lesser extent, meropenem, whereas ertapenem, cefotaxime and colistin remained unaffected. Since imipenem primarily enters *P. aeruginosa* through OprD, whereas meropenem relies on this pathway to a lesser extent, this susceptibility profile strongly suggested that PtrA targets OprD-dependent permeability rather than overall outer-membrane integrity. Importantly, PtrA-mediated resistance was fully retained in the Δ*czcRS* Δ*copRS* mutant, demonstrating that PtrA acts independently of the transcriptional pathways that repress *oprD* during Zn and Cu stress (Perron et al. 2004; Caille et al. 2007).

**Table 1.**
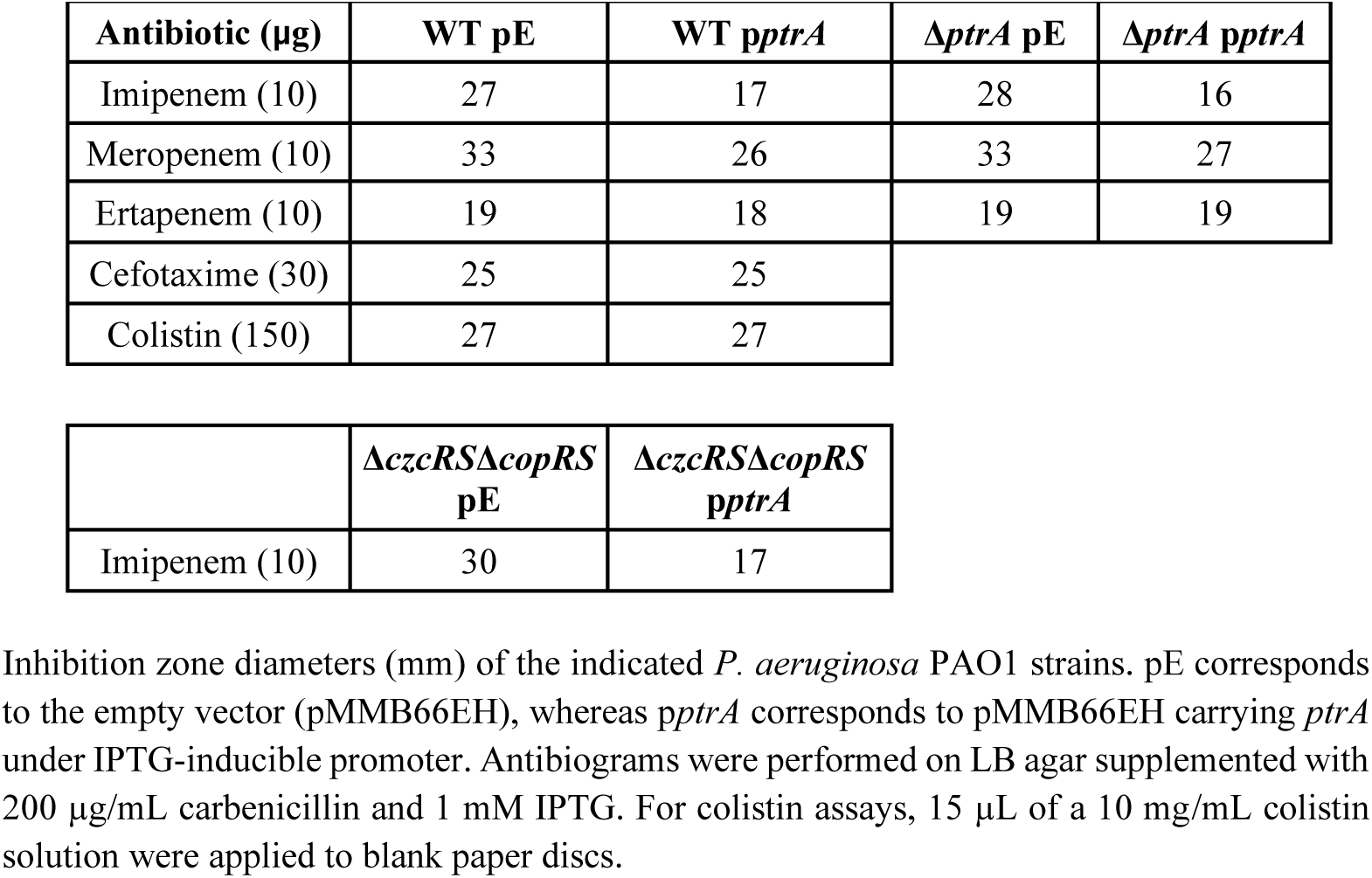
Antibiotic susceptibility profile following PtrA overexpression.

| Antibiotic (μg) | WT pE | WT *pptrA* | Δ*ptrA* pE | Δ*ptrA* *pptrA* |
| --- | --- | --- | --- | --- |
| Imipenem (10) | 27 | 17 | 28 | 16 |
| Meropenem (10) | 33 | 26 | 33 | 27 |
| Ertapenem (10) | 19 | 18 | 19 | 19 |
| Cefotaxime (30) | 25 | 25 |  |  |
| Colistin (150) | 27 | 27 |  |  |

| | Δ*czcRS*Δ*copRS*<br>pE | Δ*czcRS*Δ*copRS*<br>*pptrA* |
| --- | --- | --- |
| Imipenem (10) | 30 | 17 |

Inhibition zone diameters (mm) of the indicated *P. aeruginosa* PAO1 strains. pE corresponds to the empty vector (pMMB66EH), whereas *pptrA* corresponds to pMMB66EH carrying *ptrA* under IPTG-inducible promoter. Antibigrams were performed on LB agar supplemented with 200 μg/mL carbenicillin and 1 mM IPTG. For colistin assays, 15 μL of a 10 mg/mL colistin solution were applied to blank paper discs.

### PtrA primarily targets OprD-dependent carbapenem permeability

Because OprD is the major entry pathway for imipenem, we first examined whether PtrA-mediated resistance depended on this porin. PtrA overexpression increased the imipenem MIC of the WT strain eightfold, whereas only a modest twofold increase was observed in the Δ*oprD* mutant (Fig. 2A). These observations indicate that OprD is the major determinant of the PtrA-dependent phenotype and suggest that PtrA primarily affects OprD-mediated permeability.

**Figure 2.**
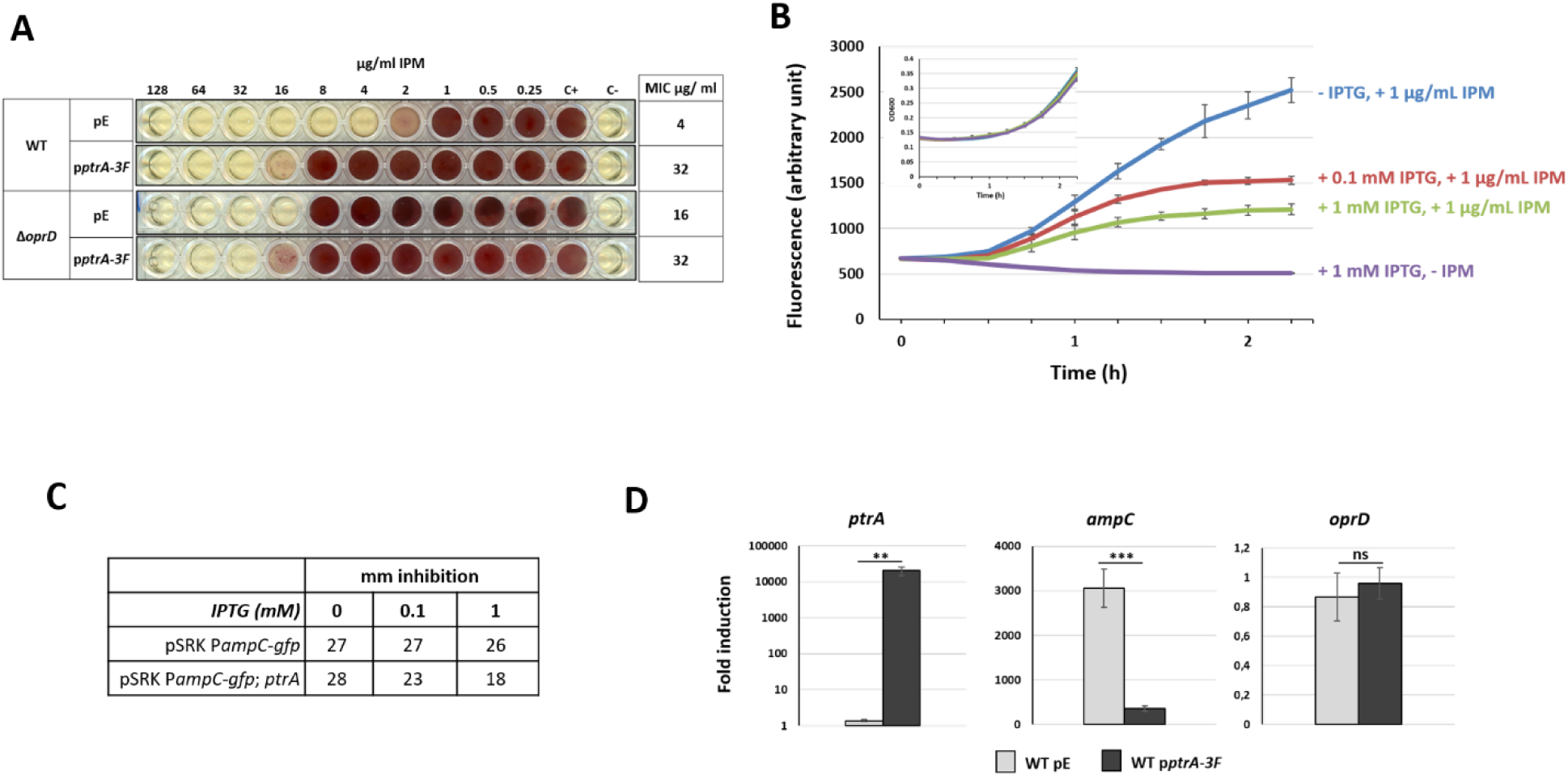
PtrA reduces OprD-dependent carbapenem permeability independently of *oprD* transcription. **(A)** PtrA-mediated resistance is primarily OprD dependent. WT and Δ*oprD* strains carrying either the empty vector (pMMB66EH) or the PtrA-3F expression plasmid were pre-induced for 3 h with 1 mM IPTG before determination of imipenem MICs by broth microdilution using two-fold serial dilutions of imipenem. Growth was visualized after 24 h using INT. MIC values are indicated on the right. **(B)** GFP fluorescence of PAO1 carrying the *PampC*-*gfp* reporter in the absence or presence of imipenem, GFP fluorescence of PAO1 carrying the P*ampC*-GFP reporter with increasing induction of *ptrA* expression (0, 0.1 or 1 mM IPTG), in the absence or presence of imipenem. Imipenem strongly induced GFP expression in the control strain, whereas induction of *ptrA* resulted in a dose-dependent reduction in reporter activity. Data represent the mean ± SD of three independent biological replicates. Bacterial growth (OD600) measured under the corresponding experimental conditions is shown in the inset. **(C)** Representative disk diffusion assays of the WT strain carrying either the P*ampC*-*gfp* reporter plasmid alone (pSRK-P*ampC*-*gfp*) or the same plasmid additionally carrying IPTG-inducible *ptrA* (pSRK-P*ampC*-*gfp*; *ptrA*), in the presence of increasing IPTG concentrations (0, 0.1 and 1 mM). Inhibition zone diameters (mm) are indicated. **(D)** qRT-PCR analysis of *ampC*, *oprD*, and *ptrA* transcript levels in WT strains carrying either the empty vector (pE) or the IPTG-inducible *ptrA*-3F plasmid (p*ptrA*-3F). Cells were grown for 3 h in the presence of 1 mM IPTG before exposure to imipenem for 1 h. Transcript levels were compared to those of the WT strain carrying the empty vector (WT pE) prior to imipenem exposure, which was arbitrarily set to 1. Data represent the mean ± SD of three independent biological replicates. Statistical significance was assessed using two-tailed unpaired Student’s *t*-tests. ns, *P* > 0.05; \**P* ≤ 0.05; \*\**P* ≤ 0.01; \*\*\**P* ≤ 0.001; \*\*\*\**P* ≤ 0.0001.

If PtrA modulates OprD function, intracellular exposure to imipenem should decrease despite unchanged OprD abundance. To test this prediction, we constructed a reporter in which the *ampC* promoter was fused to *gfp* (P*ampC*-*gfp*). Although AmpC is not a carbapenemase, *ampC* is strongly induced by imipenem and therefore provides a sensitive proxy for intracellular antibiotic exposure (Bagge et al. 2004; Freed and Hanson Nancy 2024). As expected, imipenem strongly induced GFP expression in the control strain (Fig. 2B). In contrast, overexpression of *ptrA* resulted in a marked and dose-dependent reduction in reporter activity, with maximal inhibition observed at 1 mM IPTG. These observations indicate that PtrA reduces intracellular imipenem exposure, consistent with reduced OprD-dependent permeability rather than altered antibiotic detoxification. This functional phenotype was independently confirmed by disk diffusion assays performed under identical induction conditions (Fig. 2C). Increasing PtrA expression progressively reduced the diameter of the imipenem inhibition zone, whereas IPTG had no effect in cells carrying the P*ampC*-*gfp* reporter alone. To validate that the GFP reporter faithfully reflected the endogenous transcriptional response, we quantified *ampC*, *oprD* and *ptrA* transcript levels by qRT-PCR. Cells were pre-induced with IPTG for 3 h before exposure to imipenem for 1 h (Fig. 2D). Consistent with the reporter assay, *ampC* induction was markedly reduced following PtrA expression, whereas *oprD* transcript levels remained unchanged. As expected, *ptrA* expression was strongly induced under these conditions. Collectively, these independent approaches demonstrate that PtrA reduces OprD-dependent carbapenem permeability independently of *oprD* transcriptional regulation.

### PtrA associates with OprD-containing membrane complexes

Having established that PtrA-mediated resistance is primarily OprD dependent, we next asked whether PtrA physically associates with this porin. Having established that PtrA-mediated resistance is primarily OprD dependent, we next investigated the subcellular localization of PtrA using the IPTG-inducible *ptrA*-3F construct. Cell fractionation revealed that, in addition to its expected periplasmic localization, PtrA-3F was consistently detected in the membrane fraction in the WT background (Fig. 3A). A similar distribution was observed in the Δ*opdP* mutant, indicating that deletion of *opdP* does not affect PtrA membrane association. In contrast, the membrane-associated pool was absent in the Δ*oprD* background, in which PtrA-3F was recovered exclusively in the periplasmic fraction, indicating that OprD is required for PtrA membrane association.

**Figure 3.**
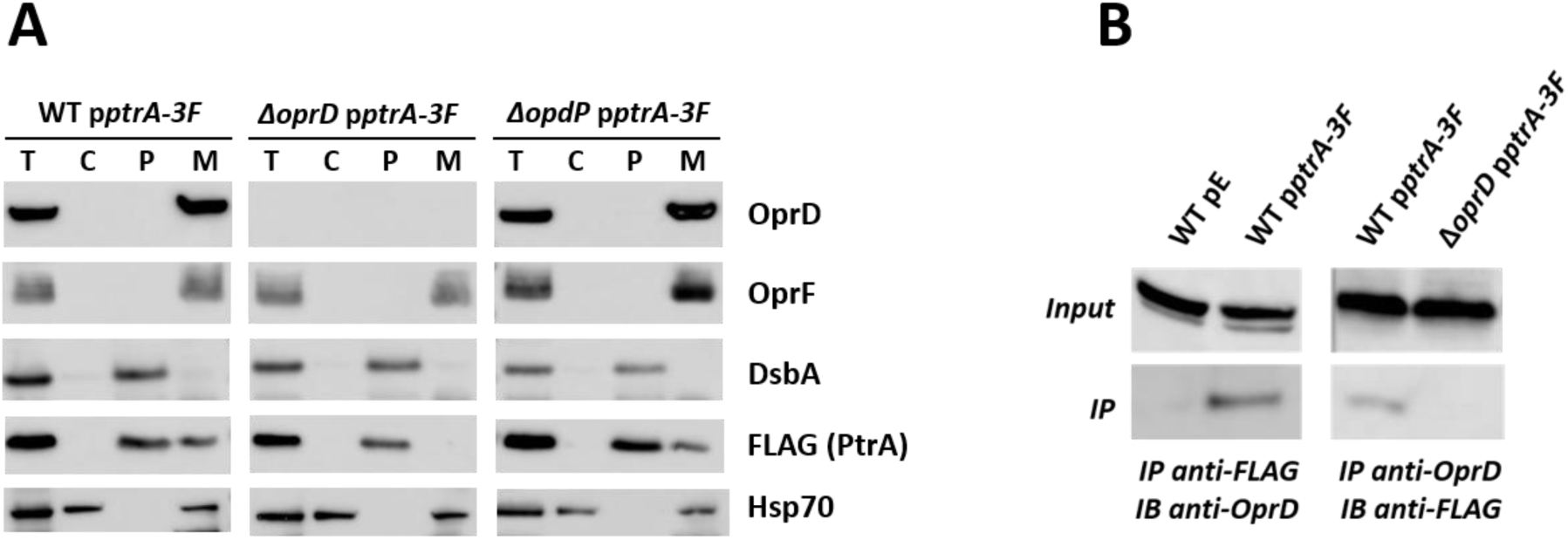
PtrA associates with OprD-containing membrane complexes. **(A)** Subcellular fractionation of WT, Δ*oprD* and Δ*opdP* strains expressing PtrA-3F, followed by immunoblot analysis. Total protein extracts (T), cytoplasmic (C), periplasmic (P) and membrane (M) fractions were probed with the indicated antibodies. OprD and OprF served as outer-membrane markers, DsbA as a periplasmic marker, and Hsp70 as a cytoplasmic marker. As previously reported, Hsp70 was also detected in the membrane fraction (el Yaagoubi et al. 1994). PtrA-3F was detected in both the periplasmic and membrane fractions of the WT and Δ*opdP* strains, whereas membrane localization was lost in the Δ*oprD* mutant. **(B)** Reciprocal co-immunoprecipitation analysis of PtrA-3F and OprD. Total protein extracts (Input) and immunoprecipitated fractions (IP) obtained using anti-FLAG or anti-OprD antibodies were analysed by immunoblotting (IB) with the indicated antibodies. OprD co-immunoprecipitated with PtrA-3F following anti-FLAG immunoprecipitation, and PtrA-3F was reciprocally recovered following anti-OprD immunoprecipitation. No corresponding signals were detected in control strains lacking either PtrA-3F or OprD.

These observations suggested a physical association between PtrA and OprD. Consistent with this hypothesis, immunoprecipitation of PtrA-3F specifically co-purified OprD, whereas no OprD signal was detected in the control strains (Fig. 3B). Reciprocally, immunoprecipitation with anti-OprD antibodies recovered PtrA-3F only in the WT PtrA-3F background. The reciprocal recovery of both proteins strongly supports a specific association between PtrA and OprD. Together, these findings demonstrate that PtrA associates with OprD-containing membrane complexes and that OprD is required for PtrA association with the outer membrane.

### Structural modelling predicts a periplasmic interface between PtrA and OprD

Having established that PtrA associates with OprD-containing membrane complexes, we next asked whether structural modelling could provide insight into the molecular basis of this interaction. AlphaFold-Multimer (v3) consistently predicted a stable complex between PtrA and the periplasmic face of OprD (Fig. 4). The highest-ranked model showed high overall confidence (pLDDT = 96.1, pTM = 0.93, ipTM = 0.896) and positioned PtrA at the periplasmic vestibule of the OprD β-barrel. Rather than interacting with the extracellular loops, PtrA was predicted to contact the periplasmic entrance of the channel consistent with a direct interaction and providing a plausible structural basis for transient modulation of porin permeability without requiring OprD degradation. The C-terminal region of PtrA remains outside the predicted interaction interface, consistent with the retained activity of the C-terminally 3×FLAG-tagged construct. Although AlphaFold predictions do not demonstrate the interaction experimentally, they provide a mechanistic framework that is fully consistent with the biochemical and functional observations described above.

**Figure 4.**
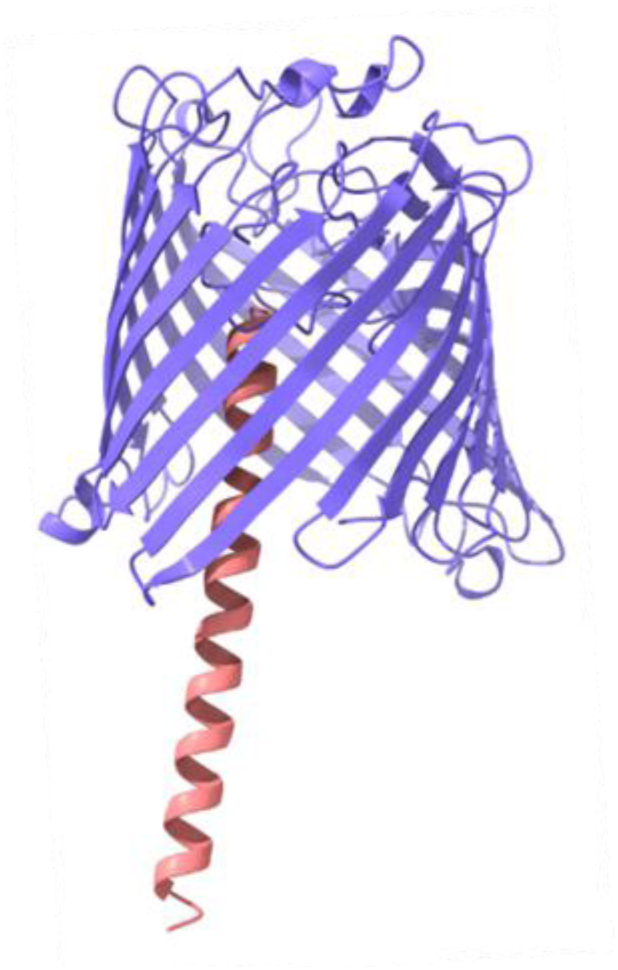
AlphaFold-Multimer prediction of the PtrA–OprD complex. Predicted interaction between the periplasmic protein PtrA (pink) and the outer-membrane porin OprD (violet) generated using AlphaFold-Multimer (v3). The highest-confidence model (pLDDT = 96.1, pTM = 0.93, ipTM = 0.896) positions PtrA at the periplasmic vestibule of the OprD β-barrel, defining a periplasmic interface between the two proteins. This predicted arrangement is compatible with transient modulation of OprD permeability without requiring porin degradation. The C-terminal region of PtrA remains exposed outside the predicted interface, consistent with the retained activity of the C-terminally 3×FLAG-tagged PtrA construct used throughout this study.

### Copper uncouples rapid periplasmic regulation from transcriptional repression of *oprD*

To determine how PtrA-mediated regulation is coordinated with the well-established CzcRS-dependent transcriptional repression of *oprD*, we monitored the temporal accumulation of PtrA, CzcR and OprD during Zn and Cu stress. For this purpose, a functional PtrA-3FLAG fusion expressed from its native promoter was integrated into the chromosomal *attB* site of the Δ*ptrA* mutant. Zn directly activates the CzcRS signalling pathway, resulting in the simultaneous induction of PtrA and transcriptional repression of *oprD*. In contrast, copper initially activates CopRS, which induces *ptrA* but activates CzcRS only at higher Cu concentrations through regulatory cross-talk (Perron et al. 2004; Caille et al. 2007). This regulatory hierarchy provides a unique experimental system to separate PtrA accumulation from CzcRS-dependent repression of *oprD* and to determine whether PtrA-mediated permeability control can occur independently of transcriptional regulation.

Exposure to 0.5 mM ZnCl₂ induced rapid accumulation of both PtrA and CzcR, whereas OprD abundance progressively declined over time (Fig. 5A). Because CzcR represses *oprD* transcription, the delayed loss of OprD is consistent with transcriptional repression followed by progressive depletion of pre-existing OprD from the outer membrane. Thus, PtrA accumulates before OprD disappears, creating a temporal window during which permeability can be regulated independently of porin abundance. In contrast, 0.5 mM CuCl₂ induced PtrA in the absence of detectable CzcR, while OprD abundance remained unchanged (Fig. 5B). Thus, copper experimentally separates PtrA accumulation from CzcRS-mediated repression of *oprD*. Increasing the Cu concentration to 2 mM induced both PtrA and CzcR and resulted in progressive loss of OprD (Fig. 5C), thereby reproducing the Zn response. Together, these experiments demonstrate that rapid periplasmic regulation by PtrA and transcriptional repression of *oprD* represent two mechanistically distinct regulatory layers that can be experimentally uncoupled using copper.

**Figure 5.**
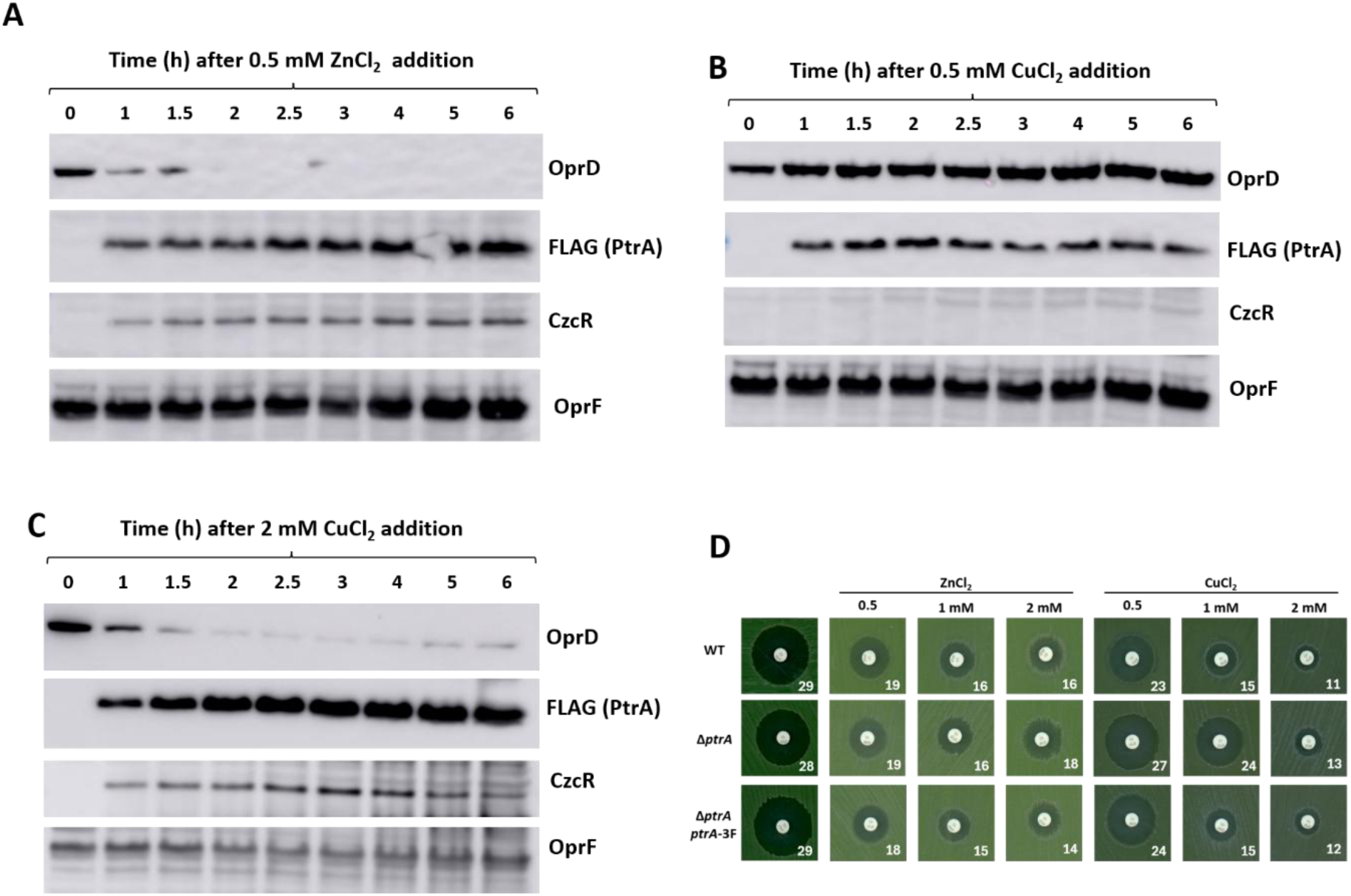
Differential regulation of PtrA and OprD during Zn and Cu stress. **(A–C)** Western blot analysis of PtrA, CzcR, OprD and OprF following exposure of *ΔptrA*::*ptrA-3F* to 0.5 mM ZnCl₂ (A), 0.5 mM CuCl₂ (B), or 2 mM CuCl₂ (C) for the indicated times. OprF was used as a loading control. **(D)** Representative imipenem disk diffusion assays performed in the absence or presence of ZnCl₂ or CuCl₂ at the indicated concentrations (0.5, 1 and 2 mM). The WT::Gm strain, the Δ*ptrA*::Gm mutant and the complemented strain (Δ*ptrA*::*ptrA-3F*) were tested under each condition. Inhibition zone diameters (mm) are indicated below each image.

These distinct regulatory states were directly reflected in imipenem susceptibility (Fig. 5D). Under Zn stress, WT, Δ*ptrA* and the complemented strain displayed comparable resistance, consistent with CzcRS-mediated repression of *oprD* in all three backgrounds. By contrast, under 0.5 mM CuCl₂, only PtrA-expressing strains exhibited reduced susceptibility, whereas the Δ*ptrA* mutant remained sensitive. Because CzcR was not induced under these conditions, this phenotype demonstrates that PtrA alone is sufficient to transiently reduce OprD permeability independently of transcriptional repression. Together, these experiments demonstrate that PtrA provides an immediate permeability control mechanism that precedes the slower transcriptional remodeling of the outer membrane.

### A two-phase mechanism of adaptive impermeability

Collectively, these findings define a two-phase mechanism of adaptive impermeability (Fig. 6). Immediately following metal sensing, PtrA accumulates in the periplasm and associates with OprD, transiently reducing OprD permeability without altering porin abundance. In a second, slower phase, activation of CzcRS represses *oprD* transcription, leading to the progressive depletion of OprD from the outer membrane and establishing sustained impermeability.

**Figure 6.**
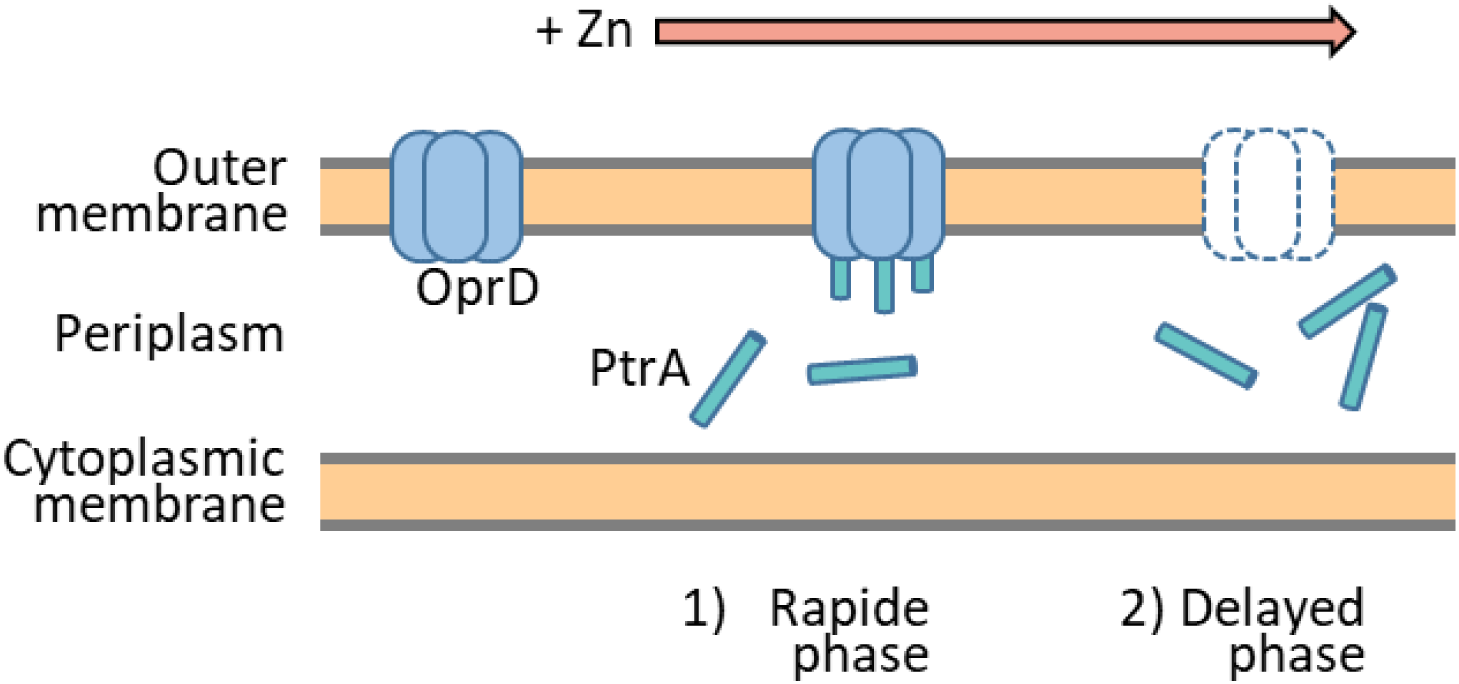
Two-phase mechanism of adaptive impermeability during metal stress. Proposed model for sequential regulation of OprD permeability following metal sensing in *P. aeruginosa*. Rapid accumulation of PtrA transiently reduces OprD permeability without altering porin abundance. Subsequent activation of CzcRS represses *oprD* transcription, progressively depleting OprD from the outer membrane and establishing sustained impermeability.

## DISCUSSION

Our work identifies PtrA as the first molecular component of a previously unrecognized mechanism of adaptive impermeability. Rather than relying exclusively on transcriptional regulation, *P. aeruginosa* rapidly modulates OprD permeability through a periplasmic regulatory layer that precedes porin depletion. By combining genetic, biochemical, and structural approaches, we show that PtrA promotes imipenem resistance without affecting OprD abundance, associates with OprD-containing membrane complexes, and likely modulates porin function during Zn or Cu stress. These findings reveal a new layer of porin regulation acting upstream of the well-characterized transcriptional repression of *oprD* by CzcRS. Reduced outer-membrane permeability is a major determinant of intrinsic and acquired antibiotic resistance in Gram-negative bacteria. Most characterized mechanisms involve transcriptional control of porin expression, alterations in porin abundance, or changes in membrane composition. In contrast, examples of proteins that directly modulate the functionality of already assembled porins remain scarce. Interactions between outer-membrane β-barrel proteins and periplasmic partners are well documented, notably within the Bam complex involved in β-barrel assembly, the Lpt machinery responsible for lipopolysaccharide transport, and other envelope-associated protein networks (Kleinschmidt 2015; Janet-Maitre et al. 2024; Yoon and Song 2024). However, these systems primarily participate in porin biogenesis or envelope maintenance rather than in the regulation of porin activity. To our knowledge, no periplasmic factor has previously been shown to regulate OprD functionality directly. PtrA therefore expands the repertoire of periplasmic proteins involved in outer-membrane adaptation and reveals a previously unrecognized mechanism linking environmental sensing to permeability control.

Interestingly, although Zn exposure triggers *oprD* repression, loss of OprD itself does not confer increased Zn or Cu tolerance (Perron et al. 2004). This observation suggests that OprD regulation during metal stress is not primarily aimed at limiting metal influx. Instead, Zn and Cu may function as environmental cues signalling entry into a broader host-associated stress response. Within macrophage phagolysosomes, elevated Zn and Cu concentrations are accompanied by reactive oxygen species, antimicrobial peptides, nutrient limitation, and multiple additional stresses (Djoko et al. 2015; Uribe-Querol and Rosales 2017; Sheldon and Skaar 2019). In this context, reducing OprD permeability may protect the bacterium from a wider range of toxic host-derived molecules rather than from metals themselves. Since OprD transports basic amino acids and small peptides, repression or functional occlusion of this channel could represent a general strategy for limiting the entry of harmful compounds encountered during infection.

This interpretation is supported by previous observations showing activation of Zn-responsive pathways during intracellular survival in macrophages (Ducret et al. 2020). Rather than acting solely as a metal-detoxification system, the CzcRS regulon may therefore function as a broader environmental sensing network that couples metal detection to rapid adaptation of the bacterial envelope. Within this framework, PtrA would provide an immediate response mechanism, allowing permeability changes to occur before transcriptional remodeling can be completed.

The fractionation, co-immunoprecipitation, and structural modeling data collectively support the formation of a PtrA-OprD complex. In the absence of OprD, PtrA loses its membrane association and remains confined to the periplasm, while reciprocal co-immunoprecipitation experiments demonstrate that both proteins are recovered within the same detergent-solubilized complex. In addition, AlphaFold-Multimer predicts a stable association between PtrA and the periplasmic vestibule of OprD. Although the precise molecular details remain to be established, the predicted architecture is compatible with a transient modulation of channel permeability. Future structural and biophysical studies will be required to determine whether PtrA directly occludes the pore or instead induces conformational changes that reduce carbapenem entry.

Although OprD accounts for the major contribution to the PtrA-dependent phenotype, a modest residual effect persisted in the Δ*oprD* mutant. Imipenem uptake in the absence of OprD has previously been reported and is thought to involve alternative entry routes (Freed and Hanson Nancy 2024). PtrA may therefore also influence one or more of these alternative permeability determinants, although their contribution to the overall phenotype appears limited. Identifying these additional determinants will provide further insight into the molecular basis of adaptive impermeability. Interestingly, PtrA paralogues are also present in several other *Pseudomonas* species, including *P. fluorescens*, *P. stutzeri* and *P. syringae* (Ducret et al. 2023). Whether these proteins fulfil a similar function remains unknown. Whether this genomic proximity reflects a conserved functional relationship between PtrA and OprD remains unknown. Whether this genomic proximity reflects a conserved functional relationship between PtrA and OprD remains unknown. Given their conservation and the widespread importance of porin-mediated permeability in environmental adaptation, it will be of particular interest to determine whether PtrA homologs also regulate outer-membrane permeability in response to metal stress or other environmental challenges, including antibiotic exposure. Such studies will establish whether the mechanism described here represents a specialized adaptation of *P. aeruginosa* or a more broadly conserved strategy among pseudomonads.

Our findings may also provide a new perspective on the previously reported role of PtrA in copper tolerance (Elsen et al. 2011). Given its periplasmic localization and its ability to associate with an outer-membrane porin, it is tempting to speculate that PtrA could similarly modulate OprC, the major copper uptake porin of *P. aeruginosa* (Bhamidimarri et al. 2021). Such a mechanism would provide a direct link between copper homeostasis and outer-membrane permeability. Testing whether PtrA associates with OprC and modulates copper uptake under copper stress will therefore represent an important direction for future studies.

More broadly, our findings introduce the concept of *adaptive impermeability*, whereby periplasmic effectors rapidly modulate porin functionality before transcriptional reprogramming and membrane remodeling occur. We propose that PtrA represents the first member of a previously unrecognized class of periplasmic porin regulators linking environmental sensing to rapid control of outer-membrane permeability. Whether adaptive impermeability represents a widespread bacterial strategy or a specialized adaptation of *P. aeruginosa* remains an important question for future studies.

## METHODS

### Bacterial strains and growth conditions

*P. aeruginosa* PAO1strain, its derivative strains and the plasmids used in this study, are listed in the Supplementary Table 1. Cultures were carried out in Luria-Bertani medium (AppliChem) and incubated at 37°C, supplemented or not with IPTG, ZnCl_2_ or CuCl_2_ at the indicated concentrations. When required, antibiotics were added to the medium at the following concentrations: 200 μg/mL carbenicillin, 50 μg/mL Gentamycin (Gm) and 50 μg/mL tetracycline (Tc) for *P. aeruginosa* or 100 μg/mL ampicillin and 15 μg/mL Tc or Gm for *E. coli*.

### Genetic manipulations

Primers used in this study and restriction enzymes for vector linearization are listed in Supplementary Table 2. PCR-amplified inserts were assembled using Gibson Assembly Master Mix (Invitrogen) according to the manufacturer’s instructions.

The pMMB66EH *ptrA* overexpression plasmid was constructed by conventional restriction-ligation cloning following BamHI/HindIII digestion of the vector and the *ptrA* insert. C-terminal 3F-tagged *ptrA* constructs in pMMB66EH, pME6182 and pSRK were generated by Gibson Assembly after vector linearization with EcoRI, BamHI and NdeI, respectively. The transcriptional reporter P*ampC*-gfp was generated by cloning the *ampC* promoter upstream of *gfp* into pBBR1 following KpnI digestion.

To investigate the effect of PtrA overexpression on a*mpC* promoter activity, the P*ampC*-*gfp* cassette was excised from pBBR1 by HindIII/XhoI digestion and ligated into either pSRK or pSRK *ptrA* digested with the same enzymes.

Deletions of *ptrA*, *oprD* and *opdP* were generated by homologous recombinations using overlap-extension PCR products cloned into the suicide vectors pEXG2 or pME3087, as indicated in Supplementary Table 2. Recombinant plasmids were amplified in *E. coli* DH5α, extracted and sequenced before introduction into *P. aeruginosa* by electroporation **(Choi et al. 2006)**. Chromosomal mutants were verified by PCR and DNA sequencing.

### Antimicrobial susceptibility assays

Minimum inhibitory concentrations (MICs) were determined according to (Wiegand et al. 2008) with minor modifications. Overnight cultures of *P. aeruginosa* strains harboring either the empty vector or the *ptrA*-3F expression plasmid were diluted to an OD_600_ of 0.1 in LB broth supplemented with carbenicillin and 1 mM IPTG and incubated for 4 h at 37 °C with shaking.

Cultures were subsequently diluted to an OD_600_ of 0.05 in fresh medium and distributed into 96-well plates containing two-fold serial dilutions of imipenem. After overnight incubation at 37 °C, p-iodonitrotetrazolium violet (INT) was added as a viability indicator, and the MIC was defined as the lowest imipenem concentration preventing visible bacterial growth.

Antibiotic susceptibility was additionally evaluated by disk diffusion assays. Bacterial suspensions were adjusted to a 0.5 McFarland standard and spread onto LB agar plates supplemented, where indicated, with the appropriate selective antibiotic and either IPTG or the indicated metal at the concentrations specified in the figures or tables legends. Antibiotic disks were applied to the agar surface, and plates were incubated overnight at 37 °C before measuring the diameters of the inhibition zones.

### RNA extraction and qRT–PCR

Overnight cultures of *P. aeruginosa* strains carrying either the empty vector or the *ptrA*-3F overexpression plasmid were diluted to an OD_600_ of 0.1 in LB medium supplemented with carbenicillin and 1 mM IPTG and incubated at 37 °C with shaking for 3 h. For experiments performed in the absence of imipenem, 0.5 ml were collected after the 3h incubation and immediately mixed with 1 ml of RNAprotect Bacteria Reagent (Qiagen). For experiments assessing *ampC* induction, imipenem was added to a final concentration of 1 µg/ ml and cultures were incubated for one additional hour before sampling.

Total RNA extraction and reverse transcription were made as previously described (Dieppois et al. 2012). Cell pellets were resuspended in 100 µl Tris EDTA buffer containing 2 mg/ ml lysozyme and incubated for 10 min at room temperature. RNA was purified using the RNeasy Mini Kit (Qiagen) according to the manufacturer’s instructions. Residual genomic DNA was removed by incubating 6 µg of total RNA with 10 U RNase-free RQ1 DNase (Promega) for 2 h at 37 °C, followed by phenol chloroform extraction and ethanol precipitation. Purified RNA was resuspended in RNase-free water.

500 ng of total RNAs were then reverse transcribed using random primers and ImProm-II Reverse Transcriptase (Promega) according to the manufacturer’s instructions. The resulting cDNAs were diluted tenfold and used as template for quantitative PCR with SYBR Select Master Mix (Thermo Fisher Scientific) and the primers listed in Supplementary Table 2. Relative transcript levels were calculated using the comparative CT method (Schmittgen and Livak 2008); with *oprF* and *ppiD* as reference genes.

### ImmunoBlotting analyses

Immunoblot analyses were performed as described previously (Ducret et al. 2016): overnight cultures of *P. aeruginosa* strains carrying either the empty vector or the *ptrA*-3F overexpression construct were diluted to an OD_600_ of 0.1 in LB medium supplemented with carbenicillin and 1 mM IPTG and grown at 37 °C with shaking. After 5 h and 24 h of growth, 1 ml of culture was collected by centrifugation, and cell pellets were resuspended in SDS–PAGE sample buffer.

For metal exposure experiments, wild-type and mutant strains carrying a gentamicin resistance cassette integrated at the chromosomal attB site, or strains expressing a chromosomally integrated *ptrA*-3F fusion under its native promoter at the attB site, were diluted to an OD_600_ of 0.1 in LB medium supplemented with 50 µg/ ml gentamicin. After 2 h of growth (T0), ZnCl_2_ or CuCl_2_ were added at the indicated concentrations, and samples were collected at the specified time points by centrifugation. Cell pellets were resuspended in SDS–PAGE sample buffer. For these experiments, 25 µg of total protein were separated by SDS–PAGE and transferred onto nitrocellulose membranes using a semi-dry transfer system (Bio-Rad) according to the manufacturer’s instructions. Membranes were incubated with antibodies against OprD and CzcR (laboratory collection; (Ducret et al. 2016), FLAG (M2 monoclonal antibody, Sigma-Aldrich) and OprF (Invitrogen). Immunoblots were visualized by chemiluminescence using the Amersham Imager 680 system.

For subcellular localization experiments, overnight cultures of WT, *ΔoprD*, and *ΔopdP* strains overexpressing the *ptrA*-3F fusion were diluted 1:100 in LB medium supplemented with carbenicillin and 1 mM IPTG and grown for 4 h at 37 °C. 1ml of culture was collected by centrifugation and resuspended in SDS–PAGE sample buffer (Total proteins). In parallel, the equivalent of 5 OD units of culture was collected by centrifugation and washed with 1 ml of 50 mM Tris–HCl (pH 7.5). Cells were resuspended in 500 µl of 200 mM MgCl_2_, 50 mM Tris–HCl (pH 7.5) and incubated for 30 min at 30 °C with gentle shaking, followed by 5 min on ice and 25 min at room temperature. Samples were centrifuged at 6,000g for 10 min at 4 °C, and the supernatant was collected as the periplasmic-enriched fraction. The resulting pellet was washed with 1 ml of 50 mM Tris–HCl (pH 7.5), resuspended in 400 µl of 50 mM Tris–HCl (pH 7.5) and subjected to ultracentrifugation at 100,000g for 45 min at 4 °C. The supernatant was collected as the cytoplasmic fraction, whereas the membrane fraction was resuspended in 400 µl of 20 mM Tris–HCl (pH 7.5) containing 0.1% SDS. For Immunoblot analysis, 25 µg of total protein and 35 µl of each subcellular fraction were separated by SDS–PAGE and analyzed as described above.

Co-immunoprecipitation assays were performed as described previously (Han et al. 2019), with modifications. Overnight cultures of *P. aeruginosa* WT carrying either the empty vector (pE) or the inducible *ptrA*-3F plasmid, and the *ΔoprD* strain carrying the inducible *ptrA*-3F plasmid, were diluted 1:100 in LB medium supplemented with 200 µg/mL carbenicillin and 1 mM IPTG and grown at 37 °C for 4 h with shaking. Cultures (100 mL) were harvested by centrifugation, washed once with 0.9% NaCl, and gently resuspended in 4 mL of IP buffer (20 mM Tris–HCl, pH 8.0, 100 mM NaCl, 0.1% NP-40, and protease inhibitor cocktail; Merck). Cells were disrupted by sonication, and lysates were clarified by centrifugation at 20,000 × *g* for 20 min at 4 °C. An aliquot of each clarified lysate was mixed with SDS–PAGE sample buffer and retained as the input fraction. For immunoprecipitation, 1.8 mL of clarified lysate was incubated with either anti-FLAG or anti-OprD antibody for 1 h 30 min at room temperature on a rotating wheel. Protein A Dynabeads (25 µL; Thermo Fisher Scientific) were then added, and incubation was continued for an additional 30 min under the same conditions. Beads were washed five times with wash buffer (20 mM Tris–HCl, pH 8.0, 150 mM NaCl, and 0.1% NP-40), and bound proteins were eluted in 100 µL of SDS–PAGE sample buffer for 10 min at room temperature with rotation. Aliquots of the input (15 µL) and immunoprecipitated fractions (40 µL) were analyzed by immunoblotting as described above.

### GFP reporter assays

GFP reporter assays were performed as previously described (Ducret et al. 2020). Overnight cultures of *P. aeruginosa* strains carrying either the P*ampC*-*gfp* reporter plasmid alone or together with the IPTG-inducible *ptrA* were diluted to an OD_600_ of 0.1 in LB medium supplemented with gentamicin and the indicated concentration of IPTG (0, 0.1 or 1 mM) and incubated at 37 °C for 3 h with shaking. Where indicated, imipenem was added to a final concentration of 1 µg/ ml. Cell growth (OD_600_) and GFP fluorescence (excitation, 485 nm; emission, 528 nm) were recorded every 15 min using a BioTek Synergy HT microplate reader. Time 0 corresponds to the addition of imipenem. For each time point, fluorescence values were divided by the corresponding OD_600_ value to obtain relative fluorescence units.

### Statistical analysis and data representation

Quantitative data are presented as the mean of three independent biological experiments, with error bars representing the standard deviation (SD). Statistical analyses were performed using two-tailed Student’s *t*-tests where indicated. Statistical significance is indicated as follows: ns, *P* > 0.05; *, *P* ≤ 0.05; **, *P* ≤ 0.01; ***, *P* ≤ 0.001; and ****, *P* ≤ 0.0001. For GFP reporter assays, fluorescence values were calculated from individual measurements normalized to the corresponding OD600 values. Data are presented as the mean ± SD of three independent biological experiments. For representative data, such as immunoblots or MIC, the images shown correspond to one representative experiment from at least three independent biological replicates.

## SUPPLEMENTARY INFORMATION

### Note on PtrA nomenclature

The designation PtrA (PA2808) in *P. aeruginosa* may lead to confusion in the literature. The name originally derived from “Pseudomonas type III repressor A” following an early report suggesting a role in regulation of the type III secretion system (Ha et al. 2004). However, subsequent work demonstrated that PA2808 is a periplasmic protein involved in copper tolerance rather than in T3SS regulation (Elsen et al. 2011). In addition, an unrelated LysR-type transcriptional regulator controlling phenazine production, chitinase expression, siderophore biosynthesis, and flagellar motility in *Pseudomonas chlororaphis* PA23 is also designated PtrA (Klaponski et al. 2014). Despite sharing the same name, these proteins are not homologous. This nomenclature overlap has already resulted in misannotation of PA2808 as a LysR-type transcriptional regulator in the literature (Zhang et al. 2022). Throughout this study, PtrA refers exclusively to the periplasmic PA2808 of *P. aeruginosa*.

**Supplementary table 1.** Strains and plasmids used in this study.

Strains and plasmids used in this study
|  | Relevant characteristic(s)<sup>a</sup> | reference or source |
| --- | --- | --- |
| ***P. aeruginosa*** |  |  |
| Wild type | PAO1 wild type | laboratory collection |
| $\Delta ptrA$ | PAO1 $\Delta PA2808$ | this study |
| WT::Gm | Gm marker integrated in the PAO1 wild type, Gm<sup>r</sup> | this study |
| $\Delta ptrA$::Gm | Gm marker integrated in the $\Delta ptrA$ mutant; Gm<sup>r</sup> | this study |
| $\Delta ptrA$::*ptrA*-3F | integrated *ptrA* with a C-terminal 3xFlag tag under its own promoter, in the $\Delta ptrA$ background; Gm<sup>r</sup> | this study |
| $\Delta oprD$ | PAO1 $\Delta PA0958$ | this study |
| $\Delta opdP$ | PAO1 $\Delta PA4501$ | this study |
| $\Delta czcR\Delta czcS$ | PAO1 $\Delta czcR\Delta czcS$ | (Caille et al. 2007) |
| $\Delta copR\Delta copS$ | PAO1 $\Delta copR\Delta copS$ | (Ducret et al. 2021) |
| **Plasmids** |  |  |
| pME3087 | Suicide plasmid, Co1E1 replicon; Tc<sup>r</sup> | (Voisard et al. 1994) |
| pEXG2 | Suicide plasmid, Co1E1 replicon; Gm<sup>r</sup> | (Rietsch et al. 2005) |
| pMMB66EH | Expressing vector carrying an IPTG-inducible promoter; Ap<sup>r</sup>, Cb<sup>r</sup> | (Fürste et al. 1986) |
| pMMB66EH-*ptrA* | pMMB66EH derivative, carrying the *ptrA* gene; Ap<sup>r</sup>, Cb<sup>r</sup> | this study |
| pMMB66EH *ptrA*-3F | pMMB66EH derivative, carrying the *ptrA* gene with a C-terminal 3xFlag Tag; Ap<sup>r</sup>, Cb<sup>r</sup> | this study |
| pSRK *PampC-gfp* | pSRK derivative plasmid, carrying the *ampC* promoter fused to the *gfp* orf; Gm<sup>r</sup> | this study |
| pSRK Plac-*ptrA* *PampC-gfp* | pSRK derivative plasmid, carrying the *pampC-gfp* transcriptional fusion and an IPTG inducible *ptrA*; Gm<sup>r</sup> | this study |

<sup>a</sup>Antibiotic resistance are indicated by r: Ap, ampicillin, Gm, gentamicin; Cb, carbenicillin.

**Supplementary Table 2.**
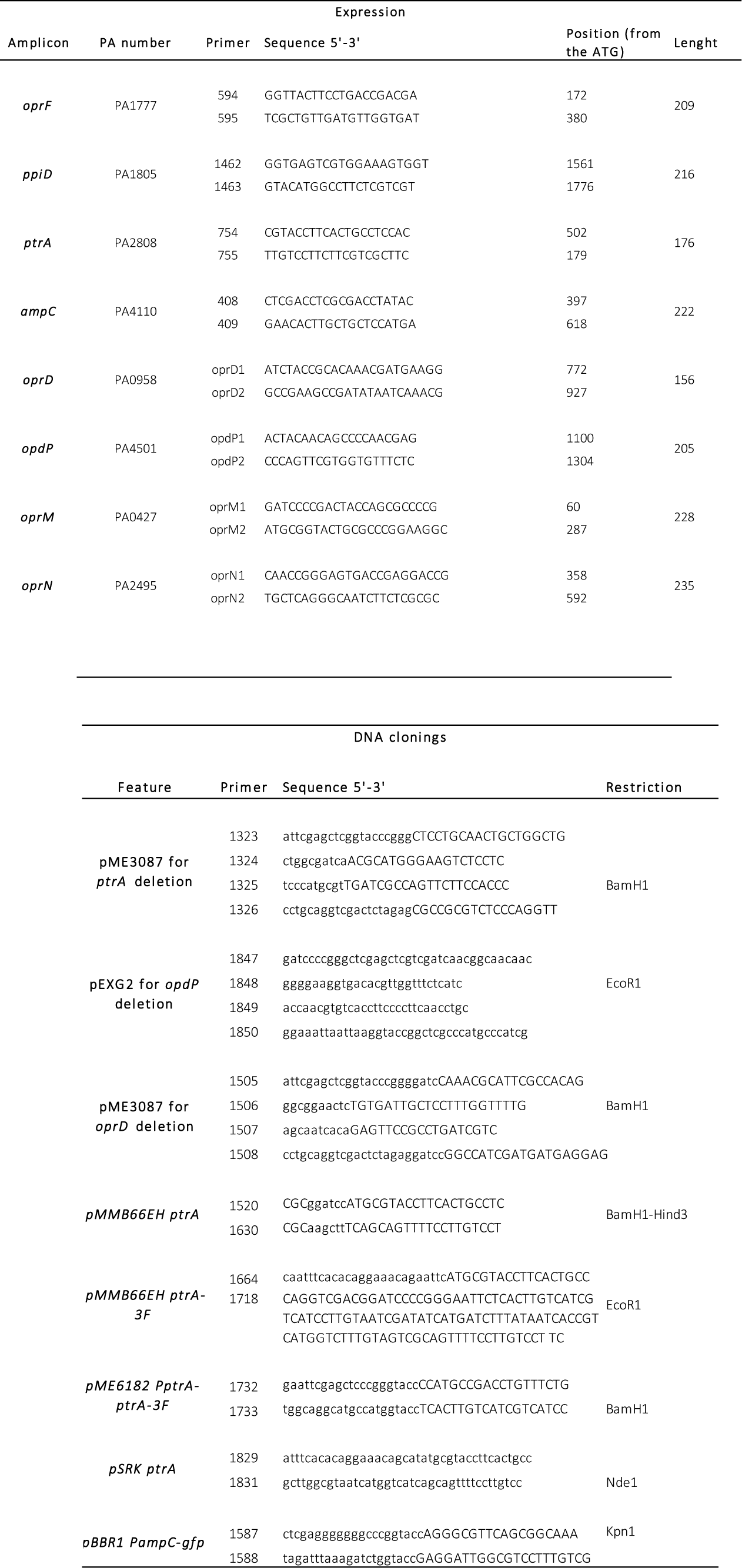
Primers used in this study.

## Notes

### Competing Interest Statement

The authors have declared no competing interest.

